# ATP-Gated Potassium Channels Contribute to Ketogenic Diet-Mediated Analgesia in Mice

**DOI:** 10.1101/2023.05.22.541799

**Authors:** Jonathan D. Enders, Sarah Thomas, Paige Lynch, Jarrid Jack, Janelle M. Ryals, Patrycja Puchalska, Peter Crawford, Douglas E. Wright

## Abstract

Chronic pain is a substantial health burden and options for treating chronic pain remain minimally effective. Ketogenic diets are emerging as well-tolerated, effective therapeutic strategies in preclinical models of chronic pain, especially diabetic neuropathy. We tested whether a ketogenic diet is antinociceptive through ketone oxidation and related activation of ATP-gated potassium (K_ATP_) channels in mice. We demonstrate that consumption of a ketogenic diet for one week reduced evoked nocifensive behaviors (licking, biting, lifting) following intraplantar injection of different noxious stimuli (methylglyoxal, cinnamaldehyde, capsaicin, or Yoda1) in mice. A ketogenic diet also decreased the expression of p-ERK, an indicator of neuronal activation in the spinal cord, following peripheral administration of these stimuli. Using a genetic mouse model with deficient ketone oxidation in peripheral sensory neurons, we demonstrate that protection against methylglyoxal-induced nociception by a ketogenic diet partially depends on ketone oxidation by peripheral neurons. Injection of tolbutamide, a K_ATP_ channel antagonist, prevented ketogenic diet-mediated antinociception following intraplantar capsaicin injection. Tolbutamide also restored the expression of spinal activation markers in ketogenic diet-fed, capsaicin-injected mice. Moreover, activation of K_ATP_ channels with the K_ATP_ channel agonist diazoxide reduced pain-like behaviors in capsaicin-injected, chow-fed mice, similar to the effects observed with a ketogenic diet. Diazoxide also reduced the number of p-ERK^+^ cells in capsaicin-injected mice. These data support a mechanism that includes neuronal ketone oxidation and activation of K_ATP_ channels to provide ketogenic diet-related analgesia. This study also identifies K_ATP_ channels as a new target to mimic the antinociceptive effects of a ketogenic diet.

## Introduction

Chronic pain negatively impacts the quality of life in 18-20% of American adults [50; 70], and emerges from various etiologies, including diabetic peripheral neuropathy. Therapeutic options for patients suffering from chronic pain are limited to medications with limited efficacy and serious complications. Many molecular mechanisms are proposed to contribute to pain in diabetic peripheral neuropathy, including inflammation [6], sensitization of transient receptor potential cation (TRP) channels TRPA1 and TRPV1 [7; 22; 32; 33; 39], and accumulation of reactive metabolites such as methylglyoxal [8; 20; 22; 32; 35]. Recent work from our group and others has identified very low-carbohydrate, ketogenic diets as a promising therapeutic strategy in preclinical models of diabetic peripheral neuropathy and chronic pain [9; 15; 19; 25; 26; 55; 56; 73]. One mechanism by which a ketogenic diet improves symptoms of diabetic peripheral neuropathy is through direct and indirect detoxification of methylglyoxal [26], a reactive metabolite that causes pain and nociception in humans and rodents by directly activating TRPA1 [2; 20; 22; 31; 32]. Methylglyoxal detoxification does not, however, completely account for the analgesic action of a ketogenic diet, as ketogenic diets improve nociception in pain models not associated with elevated methylglyoxal [9; 19; 55; 56; 73]. Here, we investigated alternative mechanisms that could contribute to the analgesic effects of a ketogenic diet.

ATP-gated potassium (K_ATP_) channels couple energetic states with membrane excitability in neuronal and non-neuronal tissues. These channels are comprised of four intracellular sulfonylurea receptor subunits (SUR1 encoded by *Abcc8*, SUR2A, and SUR2B encoded by *Abcc9*) that regulate four transmembrane inward-rectifying potassium channel subunits (Kir6.1 encoded by *Kcnj8*, Kir6.2 encoded by *Kcnj11*)[65]. These channels associate tightly with glycolytic machinery [18; 34], open when bound by Mg^2+^-ADP, and are closed by high intracellular concentrations of ATP [30; 52; 62]. Thus, under low intracellular glucose concentrations, these channels open and reduce neuronal activity in hippocampal slices [37]. Similarly, ketones inhibit glycolysis and open K_ATP_ channels, thereby reducing firing rates in slice and culture preparations from the central nervous system [44; 46]. K_ATP_ channels regulate nociception, as they are expressed in the dorsal root ganglia (DRG) and dysregulated after peripheral nerve injury [38; 45]. K_ATP_ channels also regulate opioid receptor signaling [41; 42; 53], and their genetic elimination leads to mechanical allodynia and intraepidermal fiber loss characteristic of small-fiber neuropathy [49]. However, it is unclear whether ketone oxidation is associated with the activation of K_ATP_ channels in the somatosensory nervous system or whether a ketogenic diet requires K_ATP_ channel function to mediate its analgesic effect.

We tested the hypothesis that consumption of a ketogenic diet broadly provides antinociception by activating K_ATP_ channels. As an experimental model, we fed mice a ketogenic diet for one week before receiving a single intraplantar injection of various noxious stimuli—methylglyoxal, cinnamaldehyde (a TRPA1 agonist), capsaicin (a TRPV1 agonist), or Yoda1 (a PIEZO1 agonist)—and assessed nocifensive behaviors (licking, lifting, biting of the injected paw) and markers of spinal neuron activity. Using a genetic mouse model of impaired ketone oxidation in peripheral sensory neurons and pharmacological inhibitors and activators of K_ATP_ channels, we demonstrate that a ketogenic diet prevents pain behaviors in response to a range of noxious stimuli. This antinociception depends on neuronal ketone oxidation and K_ATP_ channel activity. These results identify a novel link between ketogenic diets, ketone oxidation, nociception, and K_ATP_ channel activation.

## Materials and Methods

### Animals and Diet

All animal work was performed following review and approval by the Institutional Animal Care and Use Committee of Kansas University Medical Center. Eight-week-old C57Bl/6 mice #027 were purchased from Charles River Laboratories (Willmington, MA). Sensory Neuron Advillin-Cre Knockout of *Oxct1* (Adv-KO-SCOT) mice were bred as previously described [24]. All mice were maintained on a 12:12 light:dark cycle in the Kansas University Medical Center animal research facility. Mice were given *ad libitum* access to water and either a standard rodent chow (TD.8604; Envigo, Madison, WI; 14% fat, 32% protein, and 54% carbohydrate by kcal) or a ketogenic diet (TD.96355; Envigo, 90.5% fat, 9.2% protein, and 0.3% carbohydrate by kcal). Mice fed a ketogenic diet were provided a fresh diet every 3-4 days.

### Noxious Stimuli and Drug Administration

For systemic methylglyoxal (MGO) administration, MGO (Sigma; 40% by weight in water) was diluted in sterile saline to a working concentration of 28.8 ng/μL (pH 7.4). Control mice received an intraperitoneal (I.P.) injection of sterile saline and treated mice received 720 ng methylglyoxal I.P. in saline.

For peripheral administration in spontaneous nociception assays, MGO and cinnamaldehyde were diluted in sterile saline to working concentrations of 1.5 μg/μL (pH 7.0) and 0.65 μg/μL, respectively. Capsaicin was diluted in sterile saline with 0.5% Tween 20 to working concentrations of 0.2 μg/μL for spontaneous nociception assays and response to K_ATP_ channel blockade. Yoda1 (Tocris) was diluted in sterile saline with 5% dimethyl-sulfoxide (DMSO) to a working concentration of 355.27 ng/μL. Tolbutamide was diluted to a working concentration of 0.8 μg/μL in saline with 5% DMSO. For spontaneous nociception assay, MGO (30 μg), cinnamaldehyde (13 μg), capsaicin (4 μg), or Yoda1 (7.1054 μg) were delivered by a 20 μL subcutaneous injection to the right hind paw. Capsaicin (2 μg) and tolbutamide (8 μg) were delivered by subsequent 10 μL subcutaneous injections to the right hind paw to assess the contribution of K_ATP_ channels to antinociception by a ketogenic diet. Diazoxide (115 ng, Sigma) or levcromakalim (573 ng, MedChemExpress) were delivered in a 10 μL intraplantar injection 1 hour before capsaicin (20 μg) to assess whether K_ATP_ channel activation could prevent capsaicin-evoked nociception. These doses were based on previously published work examining the effect of these drugs on nociception [45].

### Sensory Behaviors

Sensory behavioral testing was performed at baseline and on days 1, 5, 7, 9, and 12 for animals receiving I.P. MGO injection and at baseline, 60-, and 90-minutes post-capsaicin injection for animals receiving intraplantar capsaicin and tolbutamide. Before collecting baseline data, mice were acclimated to testing areas for 30 minutes and either the mesh table for 30 minutes on at least two occasions, separated by 24 hours. Before collecting mechanical threshold data, mice were again acclimated to the testing area and mesh table for 30 minutes each. Various Von Frey microfilaments were applied to the plantar surface of the hind paw following the “up-down” method for one second. Animals were observed for either a negative or a positive response, and the mechanical withdrawal thresholds were calculated following five positive responses.

Sensory behavior in animals receiving intraplantar injections was determined by observation of spontaneous nocifensive behavior (e.g., licking, biting, lifting, and shaking the injected paw). Mice were acclimated to a clear plastic cage without bedding for 5 minutes before injection. Following intraplantar injection, mice were replaced in the cage. A blinded investigator then observed the mouse for 5 minutes following injection and recorded the total number of nocifensive events the animal displayed and the total time spent engaged in nocifensive behavior.

### Spinal Early Activation in the Dorsal Horn

To assess the response of spinal dorsal horn cells to peripheral noxious stimulation, the lumbar enlargement of the spinal cord was dissected 10 minutes following intraplantar injection and post-fixed in 4% paraformaldehyde overnight. Spinal cords were cryopreserved in 30% sucrose, frozen in Optimal Cutting Temperature Compound (Sekura Tissue-Tek) and sectioned at 30 μm on a cryostat. Sections were blocked for two hours in Superblock (ThermoFisher; Grand Island, NY), 1.5% Normal Donkey Serum, 0.5% Porcine Gelatin, and 0.5% Triton X-100 (Sigma) at room temperature. Slides were incubated overnight at 4°C with rabbit α-phospho-ERK 42/44 (1:500, Cell Signaling Technologies). Slides were next incubated with AlexaFluor-555 tagged donkey-α-rabbit secondary antibody (1:1000, Molecular Probes) for one hour and imaged with a Nikon Eclipse 90i or a Nikon Eclipse Ti2 inverted microscope. A blinded investigator counted the number of p-ERK^+^ cells in the dorsal horn grey matter across three to five independent sections for each animal. To be counted, the cell had to 1) reside within the spinal dorsal horn, 2) be roughly spherical and measure between 5 and 20 μm in diameter, and 3) display an increase in p-ERK fluorescence over background levels. The average count for each mouse was used for statistical analyses.

### Statistical Analyses

All statistical analyses were performed using R version 4.2.3 and packages “Rmisc”, “car”, “dplyr”, and “ggpubr”. All analyses for which data were collected over time were performed using a mixed-model analysis of variance (ANOVA) with repeated measures. All other analyses were performed using an N-way ANOVA. As appropriate, data were further analyzed *post hoc* by pairwise t-test or Tukey’s Honest Significant Difference (HSD), as indicated. Shapiro-Wilks and Levene tests confirmed assumptions of normal distribution and homogeneity of v, respectively. All data are presented as mean +/- standard error of the mean.

## Results

### A Ketogenic Diet Mediates a Broad Analgesic Effect

Mice were fed a ketogenic diet for one week before intraplantar injection of noxious stimuli (**Figure 1A**). Mice injected with methylglyoxal demonstrated an increased number of nocifensive behaviors (**Figure 1B** *left*; N-way ANOVA, methylglyoxal: p < 2.73e^-7^) and increased time engaged in those behaviors (**Figure 1B** *right*; N-way ANOVA, methylglyoxal: p < 7.67e^-7^, diet-methylglyoxal interaction: p < 0.0047). Mice fed a ketogenic diet one week prior to methylglyoxal injection displayed significantly fewer nocifensive behaviors (*number of nocifensive events*, N-way ANOVA, diet: p < 4.64e^-6^, diet-methylglyoxal interaction: p < 2.48e^-5^; *time engaged in nocifensive behaviors*, N-way ANOVA, diet: p < 0.00388, diet-methylglyoxal interaction: p < 0.0047).

**Figure 1.**
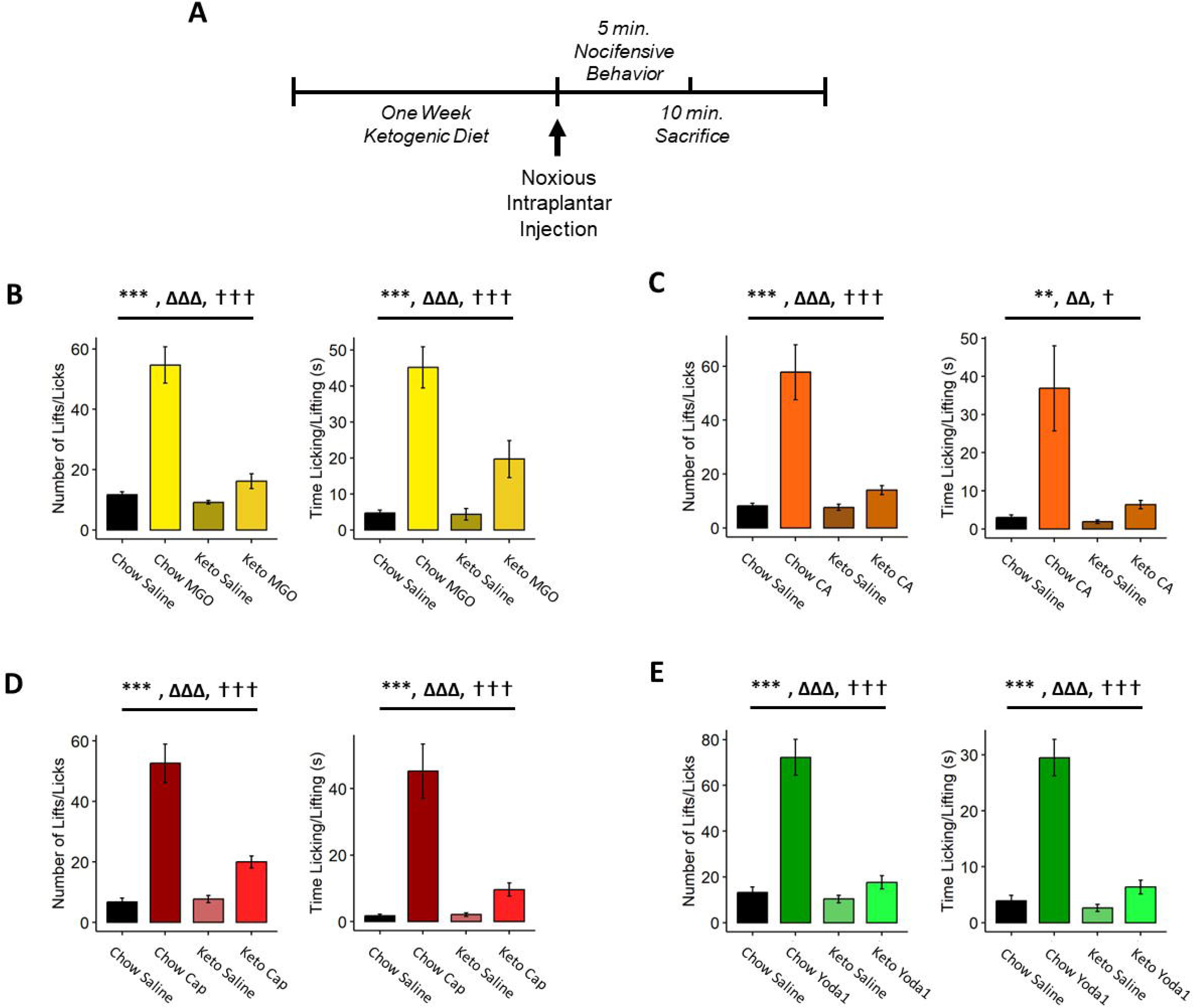
A Ketogenic Diet Provides Antinociception to Diverse Noxious Stimuli. (*A*) The experimental design for determining the effect of a ketogenic diet on nociception. Mice were fed a ketogenic diet for one week before intraplantar injection of noxious stimuli. Mice were observed for nocifensive behavior (licking, biting, lifting, etc.) for five minutes following injection. Spinal cords were collected 10 minutes following the injection of the noxious stimulus. (*B*) Chow-fed mice injected with methylglyoxal (MGO) in more nocifensive behaviors (*left*) and spent more time engaged in those behaviors (*right*), whereas mice fed a ketogenic diet were protected. Increased nocifensive behaviors were also evoked by cinnamaldehyde (CA, *C*), capsaicin (Cap, *D*), and Yoda1 (*E*) in chow-but not ketogenic diet-fed mice. (*B-E*) N-way ANOVA, ** denotes the effect of noxious stimulus: p < 0.01, *** denotes the effect of noxious stimulus: p < 0.005, ΔΔ denotes the effect of diet: p < 0.01, ΔΔΔ denotes the effect of diet: p < 0.005, ⍰ denotes the effect of stimulus-diet interaction: p < 0.05, ⍰⍰⍰ denotes the effect of stimulus-diet interaction: p < 0.005.

We postulated that the ketogenic diet-related antinociception in response to methylglyoxal-evoked nociception likely occurred too quickly to be explained by ketone bodies scavenging methylglyoxal [58]. Methylglyoxal causes pain and pain-like behaviors in humans and rodents through direct activation of TRPA1 [2; 20; 31]; thus, it is possible a ketogenic diet abrogates TRPA1-mediated nociception. To explore this possibility, we injected chow- and ketogenic diet-fed mice with cinnamaldehyde, a known TRPA1 agonist. Chow-fed mice injected with cinnamaldehyde displayed an increased number of nocifensive behaviors (**Figure 1C** *left*; N-way ANOVA, cinnamaldehyde: p < 1.05e^-5^) and engaged in nocifensive behaviors significantly longer than saline-injected mice (**Figure 1C** *right*, N-way ANOVA, cinnamaldehyde: p < 0.00199). Again, mice fed a ketogenic diet were protected from cinnamaldehyde-induced nociception (*number of nocifensive events*, N-way ANOVA, diet: p < 0.000225, diet-cinnamaldehyde interaction: p < 0.000292; *time engaged in nocifensive behaviors*, N-way ANOVA, diet: p < 0.00892, diet-cinnamaldehyde interaction: p < 0.01435).

As both methylglyoxal and cinnamaldehyde signal through TRPA1, we next tested whether ketogenic diet-mediated antinociception was effective beyond TRPA1 regulation. To this end, we used the TRPV1 agonist capsaicin. Capsaicin increased the number of nocifensive behaviors (**Figure 1D** *left*; N-way ANOVA, capsaicin: p < 3.60e^-9^) and time engaged in those behaviors (**Figure 1D** *right*; N-way ANOVA, capsaicin: p < 1.55e^-6^). Capsaicin was unable to increase the number of nocifensive events or the time engaged in nocifensive behaviors in mice fed a ketogenic diet (*number of nocifensive events*, N-way ANOVA, diet: p < 8.55e^-5^, diet-capsaicin interaction: p < 3.87e^-5^; *time engaged in nocifensive behaviors*, N-way ANOVA, diet: p < 0.000252, diet-capsaicin interaction: p < 0.000198). Together, these data suggest an antinociceptive activity of ketogenic diets that is not specific to a single TRP channel.

TRPA1 and TRPV1 channels physically interact with each other, and during this interaction activity of these channels regulate each other [61; 68]. To eliminate the possibility that the antinociceptive effect of a ketogenic diet depends on the regulation of a TRPA1-TRPV1 complex, we assessed nociceptive responses to Yoda1, a chemical activator of the mechanosensitive channel PIEZO1, following administration of a ketogenic diet. Intraplantar Yoda1 evoked a significant increase in the number of nocifensive behaviors (**Figure 1E**, *left*; N-way ANOVA, Yoda1: p < 4.03e^-08^) and the amount of time engaged in such behaviors (**Figure 1E**, *right*; N-way ANOVA, Yoda1: p < 1.17e^-8^). These behaviors were prevented in mice fed a ketogenic diet (*number of nocifensive events*, N-way ANOVA, diet: p < 5.19e^-7^, diet-Yoda1 interaction: p < 2.94^-6^; *time engaged in nocifensive behaviors*, N-way ANOVA, diet: p < 3.73e^-7^, diet-Yoda1 interaction: p < 2.27e^-6^), indicating that a ketogenic diet also reduces nociception mediated by PIEZO1 activity.

### A Ketogenic Diet Decreases Early Activation Markers in the Spinal Dorsal Horn Following Peripheral Noxious Stimuli

To measure whether a ketogenic diet affects the transmission of noxious stimuli to spinal neurons, we quantified the number of phosphorylated extracellular signal-related kinase (p-ERK) positive cells in the spinal dorsal horn ipsilateral to the intraplantar injection of a noxious stimulus. Consistent with our behavioral data, methylglyoxal increased the number of p-ERK^+^ cells in the spinal dorsal horn of chow-fed mice (**Figure 2A-B**; N-way ANOVA, methylglyoxal: p < 0.00294). Consumption of a ketogenic diet for one week prevented this increase in p-ERK^+^ cell number (N-way ANOVA, diet: p < 0.00826, diet-methylglyoxal interaction: p < 0.00306). Cinnamaldehyde increased the number of p-ERK^+^ cells in chow-fed but not ketogenic diet-fed mice (**Figure 2C-D**; N-way ANOVA, cinnamaldehyde: p < 0.02124, diet: p < 9.33e^-5^, diet-cinnamaldehyde interaction: p < 0.00499). A ketogenic diet also prevented Yoda1-dependent increases in p-ERK^+^ cell number in the spinal dorsal horn (**Figure 2E-F**; N-way ANOVA, Yoda1: p < 3.6e^-5^, diet: p < 4.41e^-6^, diet-Yoda1 interaction: p < 6.51e^-5^). These findings suggest that consumption of a ketogenic diet inhibits neuronal activation in response to noxious stimuli in the spinal dorsal horn.

**Figure 2.**
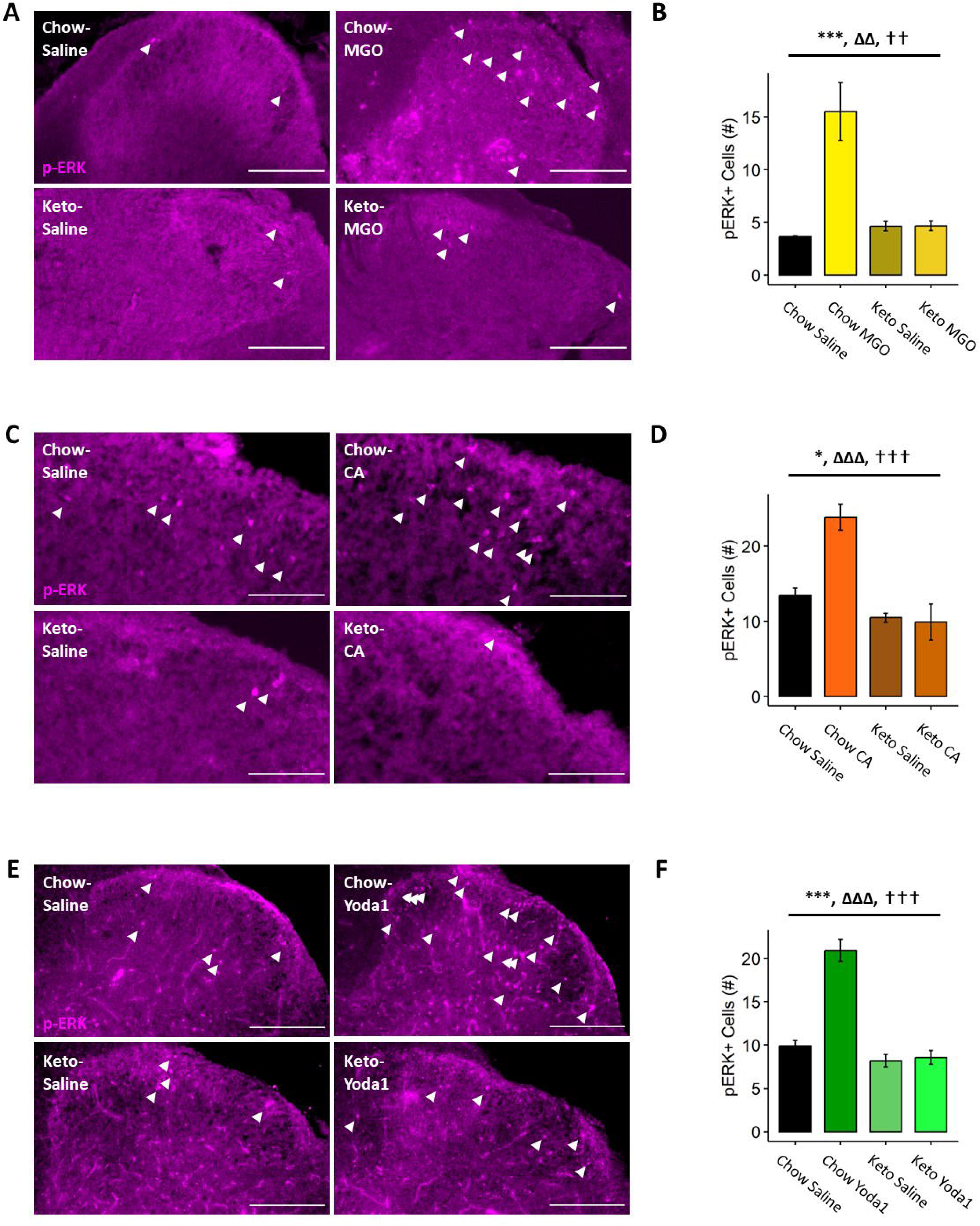
A Ketogenic Diet Prevents Noxious Stimulus-Evoked Early Activation in the Spinal Dorsal Horn. (*A-B*) Methylglyoxal increases the number of phospho-ERK (p-ERK^+^, *white arrowheads*) cells in the spinal dorsal horn within 10 minutes of intraplantar injection in chow-fed mice. Mice fed a ketogenic diet are protected from methylglyoxal-evoked early activation. (*C-D*) Cinnamaldehyde increases p-ERK^+^ cell number in the spinal dorsal horn in chow-fed mice but not mice fed a ketogenic diet. (*E-F*) Yoda1 increases p-ERK^+^ cell counts in the spinal dorsal horn of chow-fed mice within 10 minutes of injection. Mice fed a ketogenic diet are protected from Yoda1-mediated p-ERK^+^ cell number increase. (*B, D, F*) N-way ANOVA, * denotes the effect of noxious stimulus: p < 0.05, *** denotes the effect of noxious stimulus: p < 0.005, ΔΔ denotes the effect of diet: p < 0.01, ΔΔΔ denotes the effect of diet: p < 0.005, ⍰⍰ denotes the effect of stimulus-diet interaction: p < 0.01, ⍰⍰⍰ denotes the effect of stimulus-diet interaction: p < 0.005. The scale bar represents 200 μm.

### Ketone Oxidation is Required for the Analgesic Effect of a Ketogenic Diet

We next asked whether ketone oxidation as fuel in peripheral neurons was necessary to mediate the analgesic effect of a ketogenic diet. We followed the same experimental design (**Figure 1A**) but used <u>s</u>ensory <u>n</u>euron-specific <u>A</u>dvillin-<u>C</u>re <u>k</u>nockout of *<u>O</u>xct1* (Adv-KO-SCOT) mice. These mice have a tissue-specific knockout of succinyl-CoA 3-oxoacid CoA-transferase 1 (SCOT, encoded by *Oxct1*) and cannot oxidize ketone bodies for fuel in peripheral sensory neurons. As before, methylglyoxal increased the number of nociceptive events (**Figure 3A**; N-way ANOVA, methylglyoxal: p < 9.26e^-12^) and the amount of time engaged in nocifensive behavior (**Figure 3B**; N-way ANOVA, methylglyoxal: p < 6.83e^-8^) in both Adv-KO-SCOT and littermate wildtype-control mice. We also detected significant effects of genotype (*number of nocifensive events*, p < 0.033; *time engaged in nocifensive behaviors*, p < 0.01166) and genotype-methylglyoxal interaction (*number of nocifensive events*, p < 0.00739; *time engaged in nocifensive behaviors*, p < 0.00627), indicating that Adv-KO-SCOT mice may be more susceptible to methylglyoxal-evoked nociception regardless of diet. We also detected a significant effect of consuming a ketogenic diet and the interaction between consuming a ketogenic diet and methylglyoxal injection on both the number of nocifensive responses (**Figure 3A**; N-way ANOVA, diet: p < 3.34e^-5^, diet-methylglyoxal interaction: p < 2.41e^-7^) and the time engaged in those responses (**Figure 3B**; N-way ANOVA, diet: p < 0.00241, diet-methylglyoxal interaction: p < 5.17e^-6^), indicating a ketone oxidation-independent analgesic effect of a ketogenic diet.

**Figure 3.**
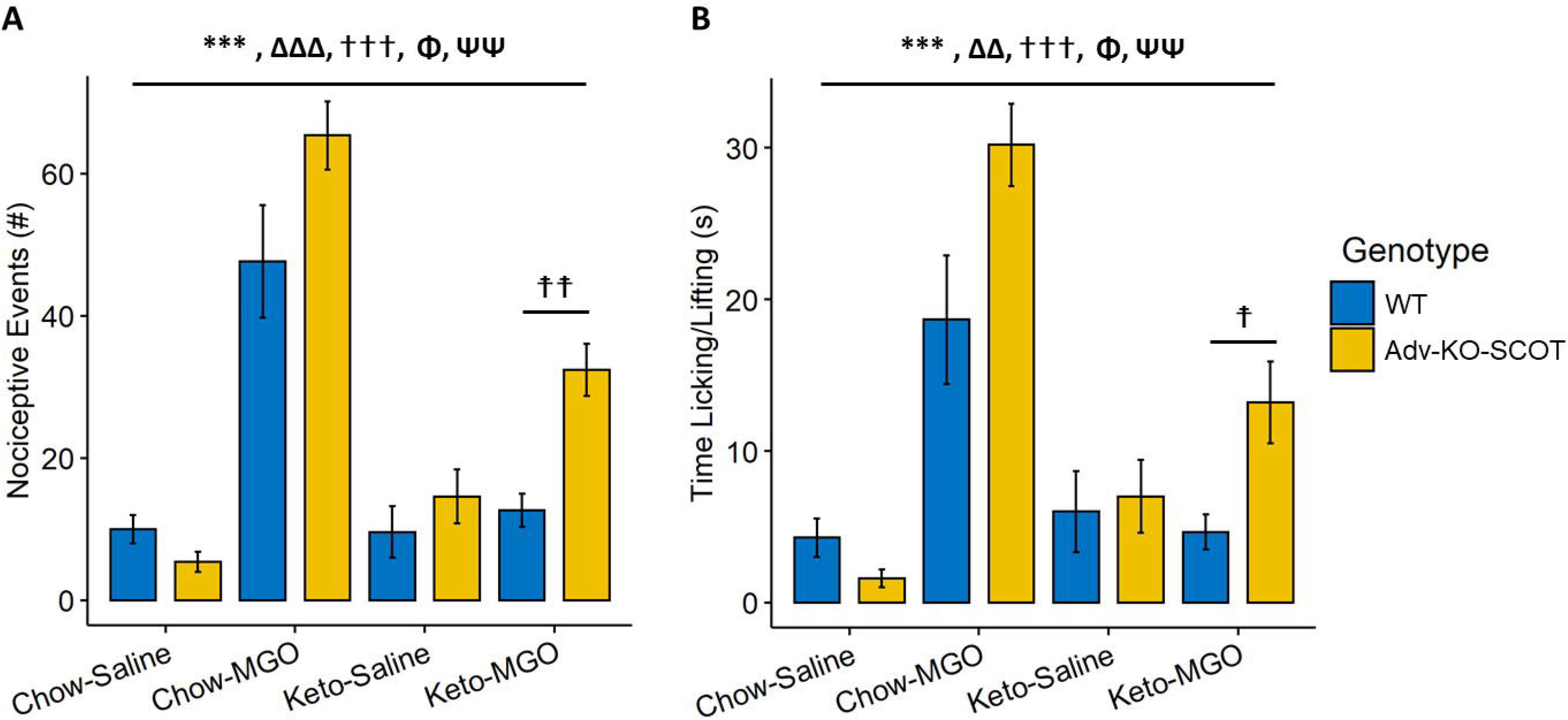
Ketone Oxidation is Required for the Full Antinociceptive Effect of a Ketogenic Diet. Wildtype (WT) and sensory neuron-specific *Advillin*-Cre knockout of *Oxct1* (Adv-KO-SCOT) mice were fed a ketogenic diet for one week before intraplantar methylglyoxal injection. Methylglyoxal increased the number of nocifensive events (*A*) and the time engaged nocifensive behaviors (*B*) in both WT and Adv-KO-SCOT mice. Consumption of a ketogenic diet reduced methylglyoxal-evoked nociception in both WT and Adv-KO-SCOT mice, though ketogenic diet-fed, methylglyoxal-injected Adv-KO-SCOT mice exhibited more nociceptive events (*A*) and engaged in nocifensive behaviors (*B*) longer than ketogenic diet-fed, methylglyoxal-injected WT mice. (*A-B*) N-way ANOVA, *** denotes the effect of noxious stimulus: p < 0.005, ΔΔ denotes the effect of diet: p < 0.01, ΔΔΔ denotes the effect of diet: p < 0.005, ⍰⍰ denotes the effect of stimulus-diet interaction: p < 0.01, ⍰⍰⍰ denotes the effect of stimulus-diet interaction: p < 0.005, Φ denotes the effect of genotype: p < 0.05, ΨΨ denotes the effect of genotype-stimulus interaction: p < 0.01. *A priori* planned comparisons, Student’s t-test ⍰ denotes the difference between WT and Adv-KO-SCOT ketogenic diet-fed, methylglyoxal injected mice: p < 0.05, ⍰⍰ denotes the difference between WT and Adv-KO-SCOT ketogenic diet-fed, methylglyoxal injected mice: p < 0.01.

Prior to data collection, we compared the responses of wildtype and Adv-KO-SCOT ketogenic diet-fed, methylglyoxal-injected mice. This analysis revealed a significant increase in the number of nocifensive responses (**Figure 3A**, planned comparison by Student’s t-test, p < 0.002692) and the time engaged in nocifensive behaviors (**Figure 3B**, planned comparison by Student’s t-test, p < 0.02995) in Adv-KO-SCOT compared to wildtype ketogenic diet-fed, methylglyoxal-injected mice. These results support a mechanism by which ketone oxidation mediates, in part, some of the protective effects of a ketogenic diet against methylglyoxal-evoked nociception.

### A Ketogenic Diet is Unable to Reduce p-ERK Expression in the Spinal Dorsal Horn in Mice Lacking the Ability to Oxidize Ketones in Peripheral Neurons

We assessed p-ERK expression in the spinal dorsal horn of ketogenic diet-fed Adv-KO-SCOT mice following methylglyoxal injection. Methylglyoxal increased the number of p-ERK^+^ cells in the spinal dorsal horn of chow-fed wildtype and Adv-KO-SCOT mice (**Figure 4**; N-way ANOVA, methylglyoxal: p < 8.74e^-9^). We detected a significant interaction between the ketogenic diet and methylglyoxal (N-way ANOVA, ketogenic diet-methylglyoxal interaction: p < 1.78e^-6^), indicating a ketone oxidation-independent effect on the number of spinal p-ERK^+^ cells following methylglyoxal injection. Prior to data analysis, we set planned comparisons *a priori* or spinal p-ERK^+^ expression between wildtype and Adv-KO-SCOT ketogenic diet-fed, methylglyoxal-injected mice. Adv-KO-SCOT ketogenic diet-fed, methylglyoxal-injected mice had significantly more p-ERK^+^ cells compared to wildtype mice (**Figure 4**, planned comparison by Student’s t-test: p < 0.000501), correlating with nociceptive behavior responses and suggesting that peripheral sensory neuron ketone oxidation is required for a ketogenic diet to reduce methylglyoxal-evoked spinal dorsal horn activity.

**Figure 4.**
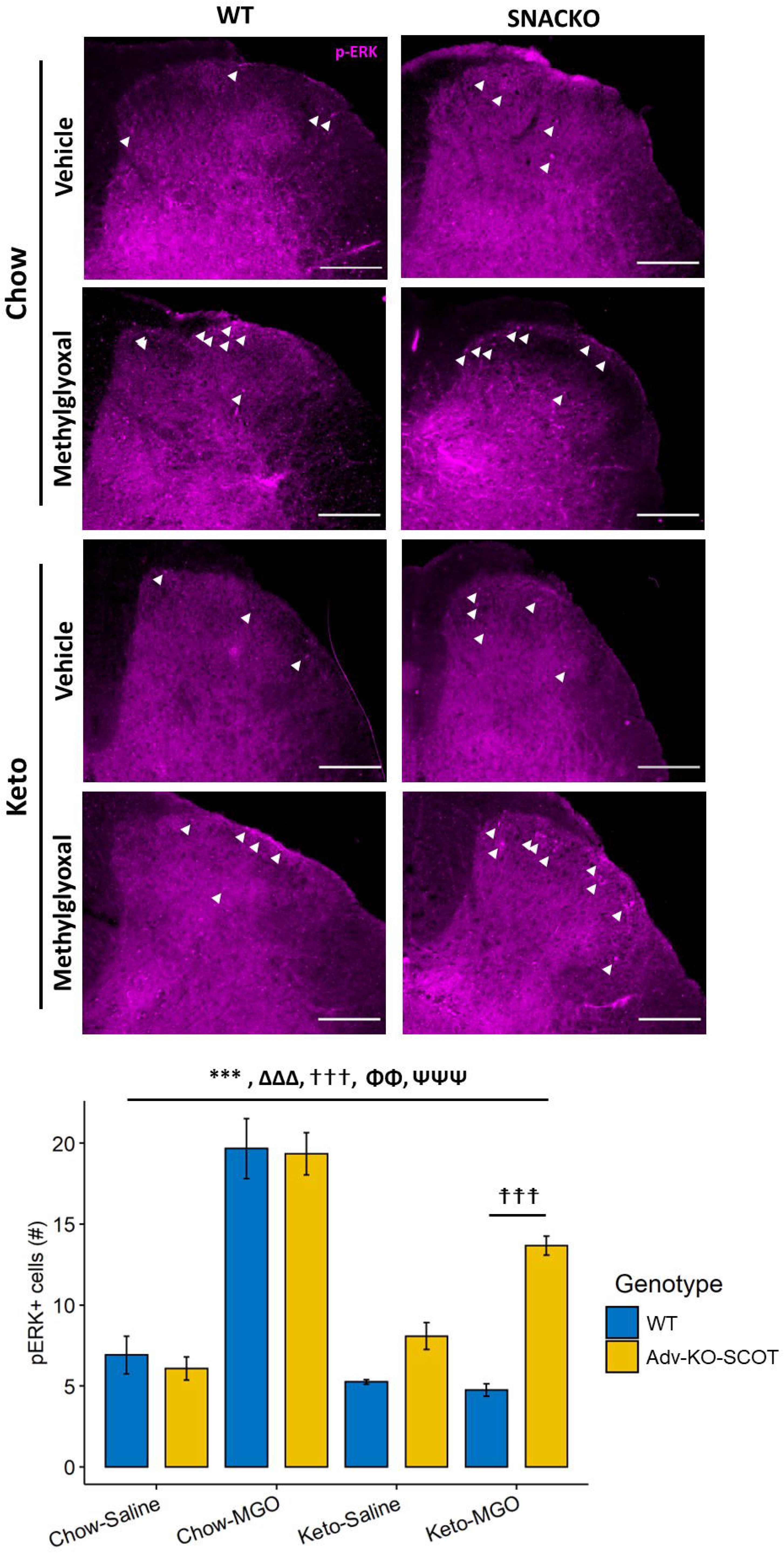
Ketone Oxidation is Required for a Ketogenic Diet to Reduce p-ERK Expression in the Spinal Dorsal Horn Following Noxious Stimuli. Intraplantar methylglyoxal injection increased the number of p-ERK^+^ cells (*white arrowheads*) in the spinal dorsal of both WT and Adv-KO-SCOT mice. Consumption of a ketogenic diet one week before methylglyoxal injection reduced the number of p-ERK^+^ cells in both genotypes; however, ketogenic diet-fed, methylglyoxal-injected Adv-KO-SCOT mice had more p-ERK^+^ cells in the spinal dorsal horn than ketogenic diet-fed, methylglyoxal-injected WT mice. N-way ANOVA, *** denotes the effect of noxious stimulus: p < 0.005, ΔΔΔ denotes the effect of diet: p < 0.005, ⍰⍰⍰ denotes the effect of stimulus-diet interaction: p < 0.005, ΦΦ denotes the effect of genotype: p < 0.01, ΨΨΨ denotes the effect of genotype-stimulus interaction: p < 0.005. *A priori* planned comparisons, Student’s t-test, ⍰⍰⍰ denotes the difference between WT and Adv-KO-SCOT ketogenic diet-fed, methylglyoxal injected mice: p < 0.005. The scale bar represents 200 μm.

### ATP-Gated Potassium (K_ATP_) Channels Are Required for Ketogenic Diet-Mediated Analgesia

To test whether the analgesic activity of a ketogenic diet requires K_ATP_ channels, we assessed the mechanical thresholds of ketogenic diet-fed mice receiving subsequent intraplantar injections of capsaicin and tolbutamide, an inhibitor of K_ATP_ channels. One hour after capsaicin injection, chow-fed mice appropriately developed mechanical allodynia, while ketogenic diet-fed mice were protected (**Figure 5A**; N-way mixed-models ANOVA with repeated measures, diet-capsaicin-time interaction: p < 0.00421; Tukey’s *post hoc* test, chow-before capsaicin-before tolbutamide compared to chow-after capsaicin-before tolbutamide: adjusted p < 1.00e^-7^, ketogenic diet-before capsaicin-before tolbutamide compared to ketogenic diet-after capsaicin-before tolbutamide: adjusted p < 1.0). In vehicle-injected mice, tolbutamide did not cause mechanical allodynia regardless of the diet consumed (Tukey’s *post hoc*, vehicle-before tolbutamide compared to vehicle-after tolbutamide: adjusted p < 0.999). Within 30 minutes of tolbutamide injection, however, ketogenic diet-fed, capsaicin-injected mice quickly developed mechanical allodynia similar to chow-fed capsaicin-injected mice (Tukey’s *post hoc*, ketogenic diet-after capsaicin-before tolbutamide compared to ketogenic diet-after capsaicin-after tolbutamide: p < 1.00e^-7^, chow-after capsaicin-before tolbutamide compared to ketogenic diet-after capsaicin-after tolbutamide: p < 1.00).

**Figure 5.**
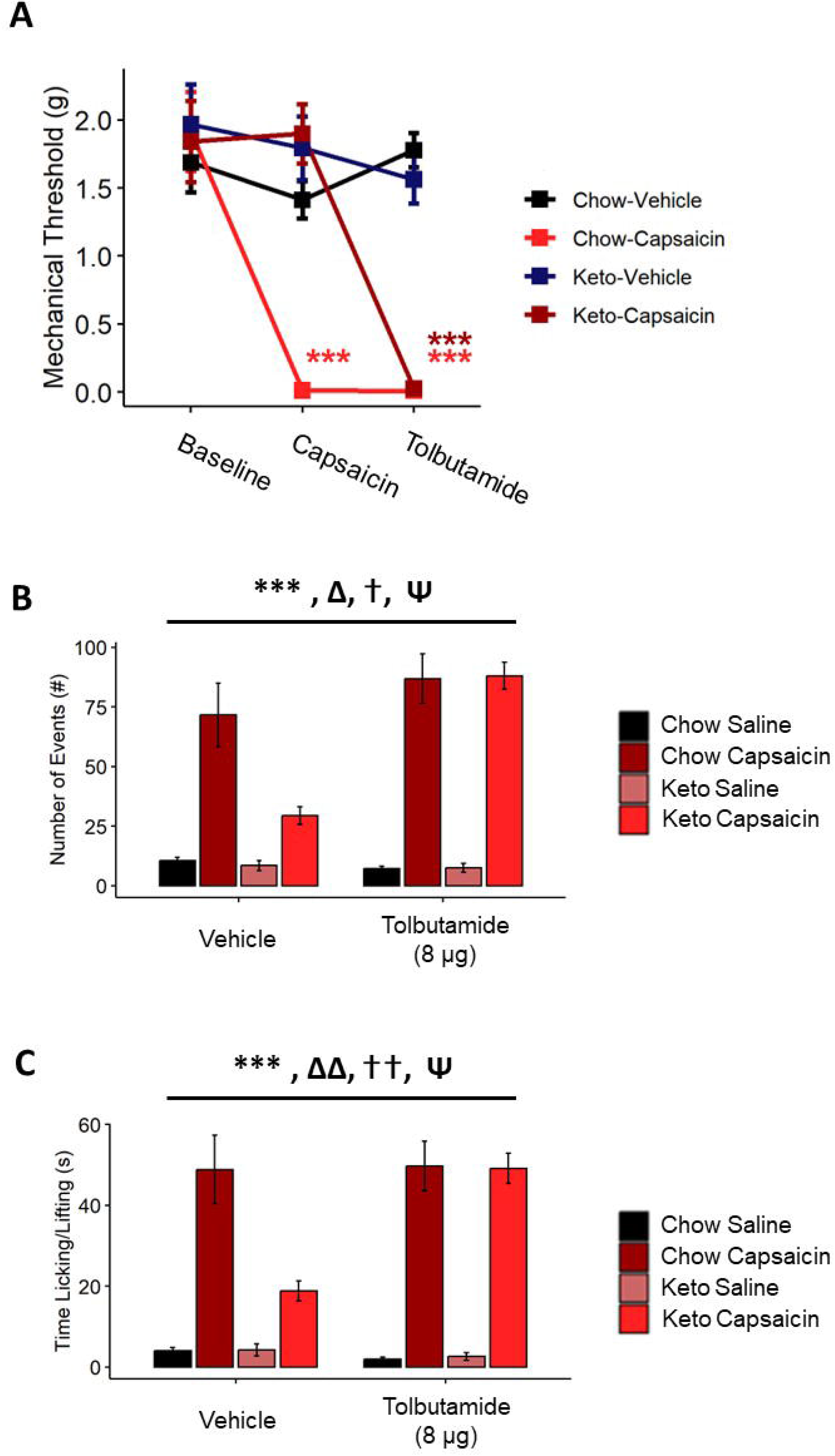
K_ATP_ Channel Activity is Required for a Ketogenic Diet to Provide an Antinociception. (*A*) Capsaicin caused mechanical allodynia in chow-fed mice one-hour following intraplantar injection. Mice fed a ketogenic diet one week before injection were protected from capsaicin-evoked mechanical allodynia. 30 minutes following ipsilateral intraplantar injection of tolbutamide, ketogenic diet-fed, capsaicin-injected mice developed mechanical allodynia. Neither chow-nor ketogenic diet-fed mice developed mechanical allodynia following tolbutamide without capsaicin. Capsaicin caused increased nociceptive events (*B*) and increased time engaged in nocifensive behaviors (*C*) in chow-fed mice but not mice fed a ketogenic diet one week before injection. Intraplantar injection of tolbutamide 30 minutes before capsaicin injection prevented protection from capsaicin-evoked nociception in ketogenic diet-fed mice. (*A*) N-way, mixed models ANOVA with Tukey’s *post hoc* test, *** indicates comparison to chow-fed, vehicle-injected: p < 0.005, color indicates the group. (*B-C*) N-way ANOVA, *** denotes the effect of noxious stimulus: p < 0.005, Δ denotes the effect of diet: p < 0.05, ⍰ denotes the effect of stimulus-diet interaction: p < 0.05, Ψ denotes the effect of stimulus-diet-tolbutamide interaction: p < 0.05.

In a separate experiment, we fed mice a ketogenic diet for one week, and tolbutamide was injected intraplantar 30 minutes before capsaicin. Consistent with our previous result, capsaicin increased the number of nocifensive behaviors (**Figure 5B**; N-way ANOVA, diet: p < 0.01107, capsaicin: p < 2e^-16^, diet-capsaicin interaction: p < 0.04189) and time engaged in nocifensive behaviors (**Figure 5C**; N-way ANOVA, diet: p < 0.00563, capsaicin: p < 2e^-16^, diet-capsaicin interaction: p < 0.0057) in chow-but not ketogenic diet-fed mice. These behavioral responses were prevented by prior injection with tolbutamide (*number of nocifensive events*, N-way ANOVA, diet-tolbutamide-capsaicin interaction: p < 0.035; *time engaged in nocifensive behaviors*, N-way ANOVA, diet-tolbutamide-capsaicin interaction: p < 0.0166), suggesting that antagonism of K_ATP_ channel activity can override the analgesic effect of a ketogenic diet.

### K_ATP_ Channels are Required for Ketogenic Diet-Mediated Reduction of p-ERK in the Spinal Dorsal Horn

We quantified p-ERK^+^ cells in the spinal dorsal horn in mice fed a ketogenic diet following tolbutamide injection. Mice receiving capsaicin had a significant increase in the number of p-ERK^+^ cells in the spinal dorsal horn compared to vehicle-injected mice (**Figure 6**; N-way ANOVA, capsaicin: p < 1.24e^-13^). This increase was prevented in mice fed a ketogenic diet (N-way ANOVA, diet: p < 1.15e^-9^, diet-capsaicin interaction: p < 4.57e^-9^). The ketogenic diet-mediated reduction in spinal p-ERK^+^ cells following capsaicin injection was prevented, however, by injection of tolbutamide (N-way ANOVA, diet-tolbutamide-capsaicin interaction: p < 0.000107). This result suggests that a ketogenic diet requires K_ATP_ channel activity to prevent peripheral noxious stimuli from reaching the spinal dorsal horn.

**Figure 6.**
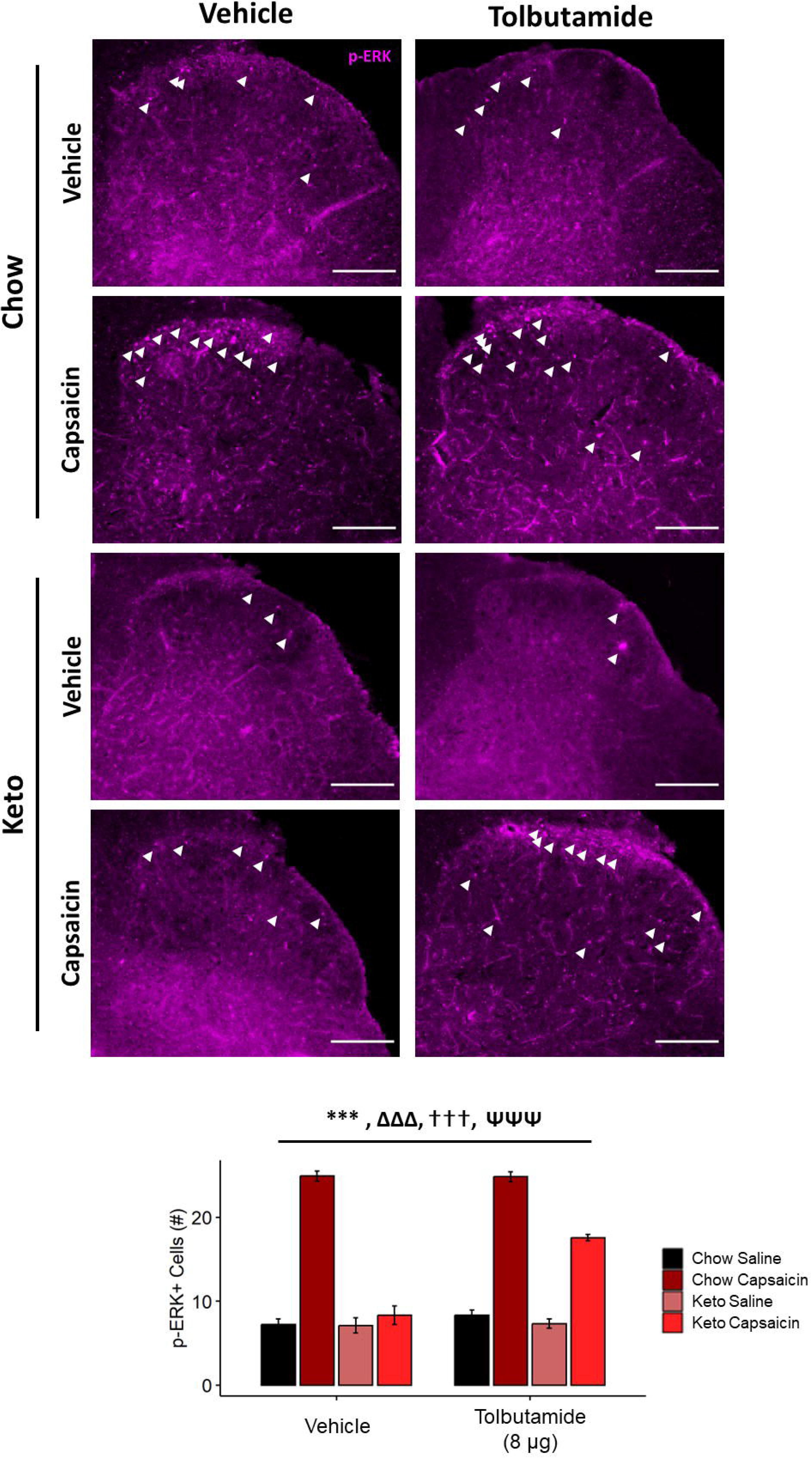
K_ATP_ Channels Are Required to Reduce p-ERK Expression Following Capsaicin Injection in Mice Fed a Ketogenic Diet. Intraplantar capsaicin injection increased the number of p-ERK^+^ cells (*white arrowheads*) in the spinal dorsal horn of chow-fed mice. Mice fed a ketogenic diet before capsaicin injection were protected from this increase in p-ERK^+^ cells. Intraplantar injection of tolbutamide 30 minutes before capsaicin injection increased the number of p-ERK^+^ cells in the spinal dorsal horn of ketogenic diet-fed mice. Tolbutamide injection did not affect the number of p-ERK^+^ cells in chow-fed capsaicin-injected mice. N-way ANOVA, *** denotes the effect of noxious stimulus: p < 0.005, ΔΔΔ denotes the effect of diet: p < 0.005, ⍰⍰⍰ denotes the effect of stimulus-diet interaction: p < 0.005, ΨΨΨ denotes the effect of stimulus-diet-tolbutamide interaction: p < 0.005.

### Peripheral Activation of SUR1-Containing K_ATP_ Channels Mediates Analgesic Effect of a Ketogenic Diet

To determine whether activation of K_ATP_ channels was sufficient to recapitulate antinociception provided by a ketogenic diet, we injected chow-fed mice with diazoxide, a broad spectrum K_ATP_ channel activator, one hour before intraplantar capsaicin injection. Mice injected with diazoxide before receiving capsaicin were protected from capsaicin-evoked nocifensive events (**Figure 7A**, N-way ANOVA, capsaicin: p < 1.14e^-8^, diazoxide: p < 0.000217, capsaicin-diazoxide interaction: p < 0.000156) and spent less time engaged in nocifensive behaviors (**Figure 7B**, N-way ANOVA, capsaicin: < 4.24e^-8^, diazoxide: p < 0.000203, capsaicin-diazoxide interaction: p < 0.000203) compared to vehicle-injected mice.

**Figure 7.**
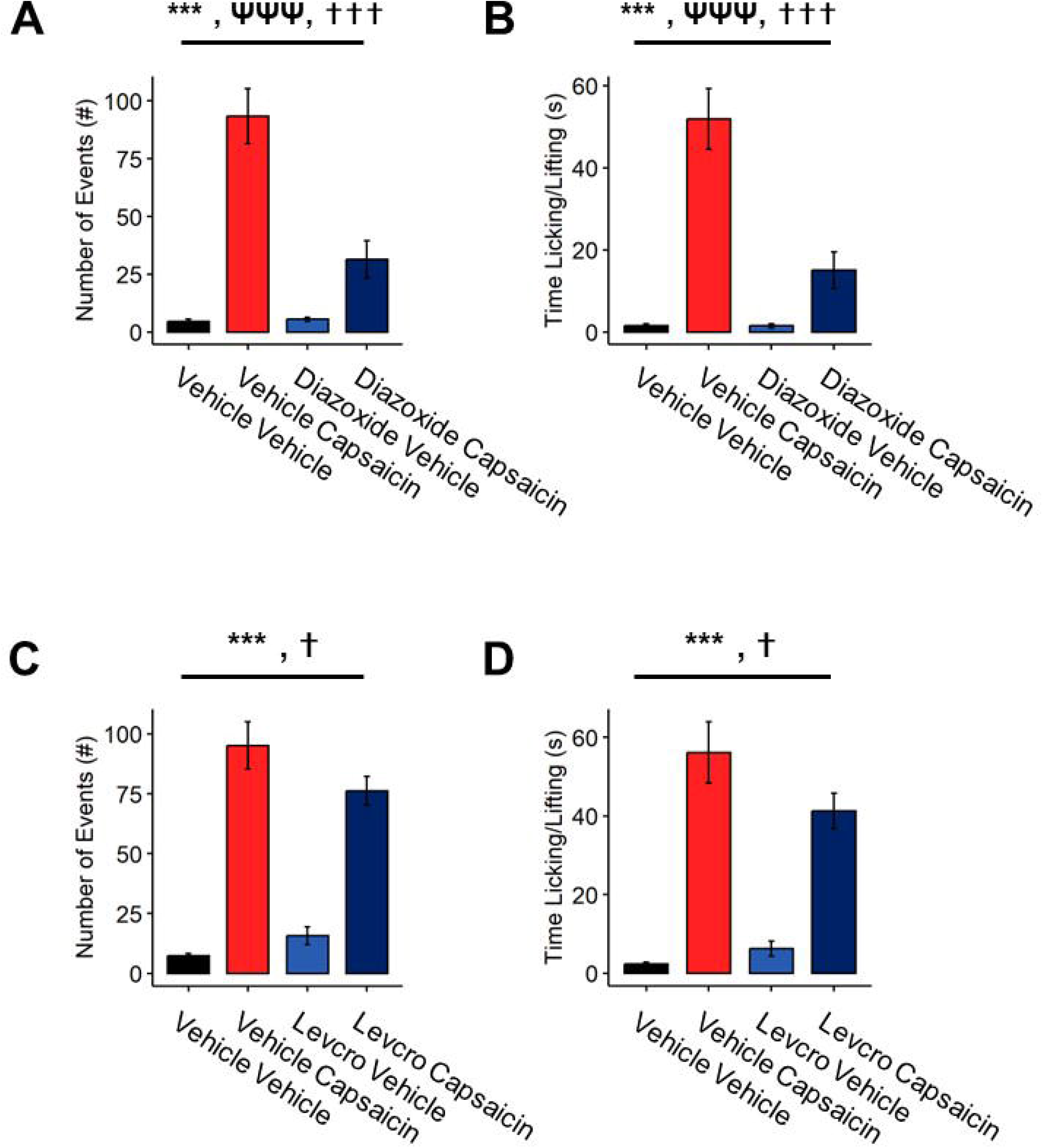
Activation of SUR1-Containing K_ATP_ Channels Mimics the Antinociceptive Effect of a Ketogenic Diet. Capsaicin increased nociceptive events (*A, C*) and increased time engaged in nocifensive behavior (*B, D*) in chow-fed mice. Intraplantar injection of diazoxide one hour before capsaicin injection was sufficient to rescue capsaicin-evoked nociception (*A-B*), while intraplantar injection of levcromakalim (levcro) offered only modest protection (*C-D*). (*A-D*) N-way ANOVA, *** denotes the effect of noxious stimulus: p < 0.005, ΨΨΨ denotes the effect of K_ATP_ channel openers: p < 0.005, ⍰ denotes the effect of stimulus-K_ATP_ openers interaction: p < 0.05, ⍰⍰⍰ denotes the effect of stimulus-K_ATP_ openers interaction: p < 0.005.

While diazoxide primarily activated K_ATP_ channels containing the SUR1 regulatory subunit, SUR2B-containing K_ATP_ channels retain some sensitivity to diazoxide [36; 47; 69]. Levcromakalim, however, specifically activates K_ATP_ channels containing SUR2A and SUR2B, but not SUR1 [59]. To determine whether there was specificity between K_ATP_ channel subunit composition and antinociception, we injected mice with levcromakalim one hour before intraplantar capsaicin injection. Mice injected with levcromakalim before capsaicin showed only modest protection from the number of capsaicin-evoked nocifensive behaviors (**Figure 7C**, N-way ANOVA capsaicin: p < 5.35e^-13^, levcromakalim: p < 0.688, capsaicin-levcromakalim interaction: p < 0.0251). Levcromakalim also modestly reduced the time engaged in capsaicin-evoked nocifensive behaviors (**Figure 7D**, N-way ANOVA capsaicin: p < 9.21e^-11^, levcromakalim: p < 0.397, capsaicin-levcromakalim interaction: p < 0.0388). While we detected a significant interaction between levcromakalim and capsaicin injection for both the number of nocifensive events and the amount of time engaged in nocifensive behavior, these behaviors were not significantly different between mice injected with vehicle and capsaicin and those injected with levcromakalim and capsaicin (number of nocifensive events: Tukey’s *post hoc*: p = 0.127; time engaged in nocifensive behaviors: Tukey’s *post hoc*: p = 0.102).

While capsaicin increased the number of p-ERK^+^ cells in the spinal dorsal horn of vehicle-injected mice (**Figure 8A-B**, N-way ANOVA, capsaicin: p < 4.95e^-5^), diazoxide significantly reduced the number of p-ERK^+^ cells (N-way ANOVA, diazoxide: p < 0.000314, capsaicin-diazoxide interaction: p < 0.003498). Prior injection with levcromakalim did not prevent increased p-ERK^+^ cell count in the spinal dorsal horn following capsaicin injection (**Figure 8C-D**; N-way ANOVA, capsaicin: p < 5.72e^-7^, levcromakalim: p = 0.569, capsaicin-levcromakalim interaction: p < 0.00684).

**Figure 8.**
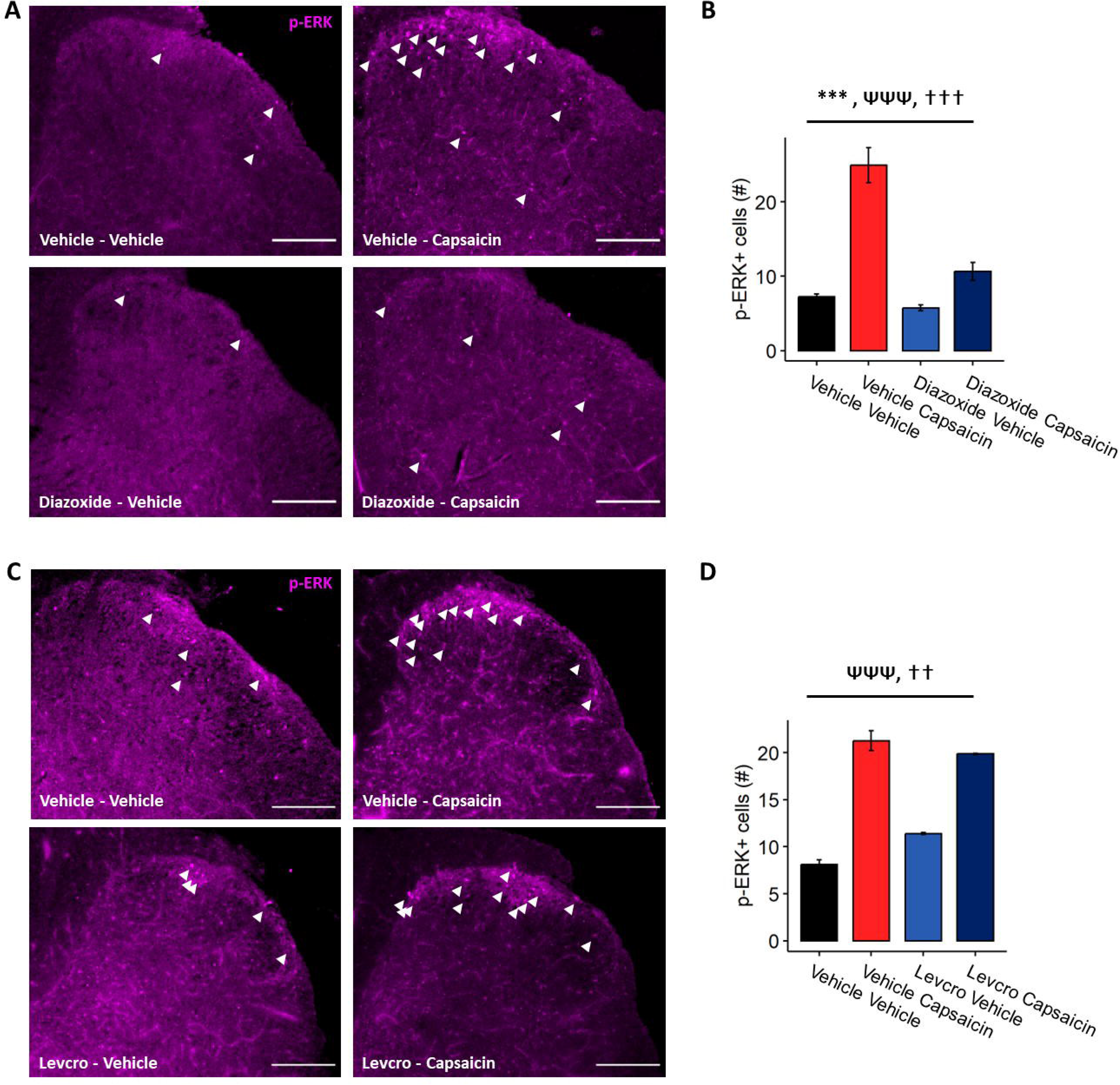
Diazoxide Reduces p-ERK^+^ Cell Number in the Spinal Dorsal Horn of Mice Receiving Intraplantar Capsaicin. (*A-D*) Intraplantar capsaicin increased the number of p-ERK^+^ cells (*A, C*, *white arrowheads*) in the spinal dorsal horn of mice. (*A-B*) Prior injection of diazoxide prevented spinal neuronal activation, whereas prior injection of levcromakalim (levcro, *C-D*) did not affect increased p-ERK^+^ cell number. (*B, C*) N-way ANOVA, *** denotes the effect of noxious stimulus: p < 0.005, ΨΨΨ denotes the effect of K_ATP_ channel openers: p < 0.005, ⍰ denotes the effect of stimulus-K_ATP_ openers interaction: p < 0.05, ⍰⍰⍰ denotes the effect of stimulus-K_ATP_ openers interaction: p < 0.005.

## Discussion

There is mounting evidence that consuming a ketogenic diet is protective against pain and pain-like behaviors in clinical and preclinical settings. Our group has previously reported that consumption of a ketogenic diet prevents and reverses mechanical allodynia in mouse models of obesity [15] and diabetic peripheral neuropathy [25]. Others have demonstrated that ketogenic diets reduce pain-like behaviors in rodent models of inflammatory pain [55; 56] and chemotherapy-induced neuropathy [73] and reduce pain in patients with migraine [9; 19] and chronic musculoskeletal pain [28]. However, the mechanisms underlying this analgesia remain poorly defined. Possible contributions include improved metabolic neuronal health, as mice fed a ketogenic diet produce fewer reactive oxygen species (ROS) in their sciatic nerve [14], and a ketogenic diet and ketone availability contribute to methylglyoxal detoxification [26; 58], It is plausible that this mechanism is important in reducing pain-like behaviors in experimental models of diabetic neuropathy [8; 22; 32; 54; 72].

In this study, we demonstrate that a ketogenic diet is broadly antinociceptive to several different noxious stimuli, including capsaicin, cinnamaldehyde, methylglyoxal, and Yoda1. As expected, intraplantar injection of these noxious agents induced nocifensive behaviors (licking, lifting, biting, etc. of the injected paw) in mice. These behaviors were diminished in mice fed a ketogenic diet for all these stimuli (**Figure 1**). Methylglyoxal-evoked nociception has been best-studied through activation of TRPA1 [2; 20; 22; 31]; however, inhibition of TRPA1 does not fully prevent nociception after methylglyoxal [5], and methylglyoxal promotes pain-like behaviors through TRPA1-independent mechanisms that include glycation of Na_V_1.8 [8], activation of the receptor for advanced glycation end products (RAGE) [67], and activation of the integrated stress response [5]. Thus, the ability of a ketogenic diet to reduce pain-like behaviors in response to cinnamaldehyde (TRPA1 agonist), capsaicin (TRPV1 agonist), and methylglyoxal suggests the antinociceptive mechanism of a ketogenic diet is not restricted to methylglyoxal detoxification [26].

TRPA1 and TRPV1 physically interact and modify each other’s activity [61; 68]. Thus, one potential antinociceptive mechanism of a ketogenic diet is regulation of the TRPA1-TRPV1 macromolecular complex. While data here do not eliminate this possibility, we demonstrate that consumption of a ketogenic diet prevents nociceptive behaviors in response to Yoda1, a chemical activator of PIEZO1. PIEZO1 is a mechanosensitive channel expressed by neuronal and non-neuronal cells [16; 40; 48; 64]. Due to the differential expression of TRPV1, TRPA1, and PIEZO1, it is unlikely that PIEZO1 activity is regulated by these other channels [64]. Since consumption of a ketogenic diet prevents PIEZO1-mediated nociception, we believe that a ketogenic diet provides a broader mechanism of antinociception than modifying these ion channels individually.

As such, we focused on the possibility of reducing the membrane excitability of nociceptors. In slice and dissociated culture recordings from the central nervous system, ketones can reduce spontaneous firing rates associated with reduced glycolytic flux and activation of K_ATP_ channels [44; 46]. K_ATP_ channels are inhibited by ATP [30; 52; 62] and associate closely with cellular glycolytic machinery [18; 34]. Others have suggested that glycolysis increases ATP concentrations near the membrane, leading to the closure of K_ATP_ channels. Ketones inhibit glycolysis [44; 46; 63], shifting the cell toward oxidative metabolism in the mitochondria and diminishing the membrane-proximal ATP pool. This combination results in the opening of K_ATP_ channels and hyperpolarization of the cell.

Relevant to pain, dysregulation of K_ATP_ channels has been reported in rodent models of neuropathy. Upregulation of K_ATP_ channel subunits occurs following spinal nerve ligation [45] and in a model of diabetic neuropathy [49]. This may represent a compensatory mechanism, as knockout of the Kir6.2 or SUR1 subunits results in mechanical and thermal hyperalgesia [45; 49] and spinal nerve ligation reduced K_ATP_ conductance in the DRG of mice following spinal nerve ligation [38]. Glibenclamide, another sulfonylurea antagonist of K_ATP_ channels, increases the resting membrane potential of uninjured DRG sensory neurons [38]. Conversely, opening K_ATP_ channels with the agonist diazoxide reduces C-fiber excitability in response to mechanical stimuli [45].

Collectively, these results suggest a mechanism in which K_ATP_ channel activity hyperpolarizes neurons to reduce their firing (**Figure 9**), consistent with the results presented here. As in previous studies, we show that mice fed a ketogenic diet were protected from methylglyoxal-evoked nociception [26]. It is important to point out that Adv-KO-SCOT mice were not fully protected from methylglyoxal-evoked nociception (**Figure 3**). These mice lack expression of *Oxct1* in peripheral sensory neurons and cannot oxidize ketone bodies as fuel. Thus, we reason that the ketogenic diet-related partial protection in these mice is due to methylglyoxal detoxification [26] and ketone oxidation. In a previous study, we reported increased circulating ketones in Adv-KO-SCOT mice fed a ketogenic diet [24], indicating that ketone bodies are present to contribute to methylglyoxal scavenging [26] and alternative mechanism of anti-nociception, including resolution of inflammation [60; 71] and induction of antioxidant response [12; 43; 66]. In the absence of ketolysis, sensory neurons in ketogenic diet-fed mice likely utilize the minimal dietary carbohydrates for glycolysis, as neurons metabolize free fatty acids poorly [23]. Glycolysis increases membrane-proximal ATP concentrations, inhibiting K_ATP_ channels and preventing the hyperpolarization of nociceptors, a point that requires further testing in diabetes models.

**Figure 9.**
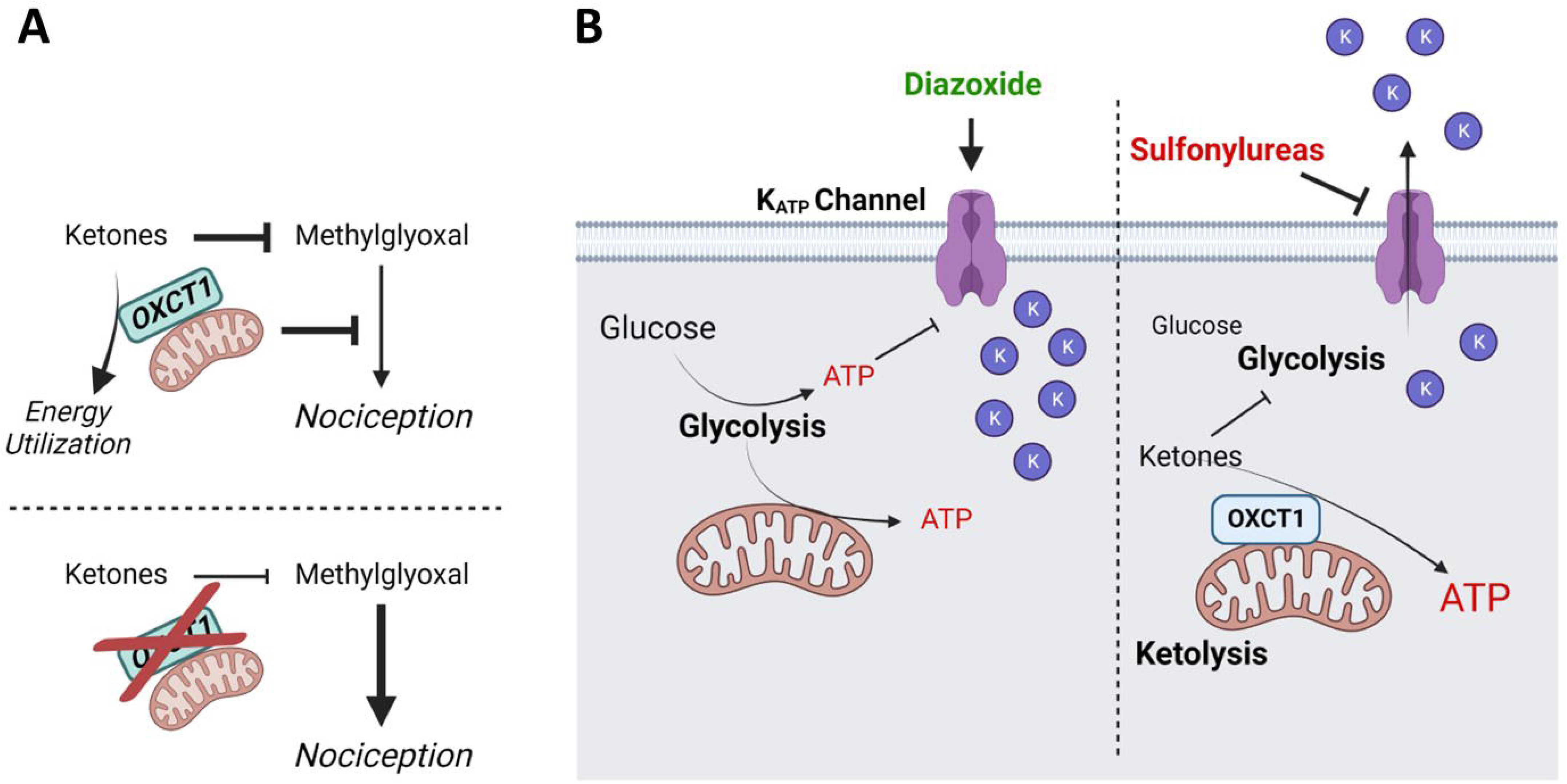
Ketone Oxidation and K_ATP_ Channel Activity are Required for the Full Analgesic Effect Provided by a Ketogenic Diet. (*A*) Ketone bodies contribute to protection from methylglyoxal-evoked nociception through direct scavenging [26] and through a second mechanism requiring *Oxct1* and ketone oxidation. (*B*) Under normoglycemic conditions (*left*), glucose is metabolized as fuel by glycolysis. As glycolytic machinery is membrane-bound and associated with K_ATP_ channels [11; 18; 27], glycolysis increases membrane-proximal concentrations of ATP and closes K_ATP_ channels. During ketosis (*right*), ketone bodies inhibit glycolysis [44] and are oxidized in the mitochondria, decreasing membrane-proximal concentrations of ATP and increasing ATP concentrations further from the membrane. This in turn allows K_ATP_ channel-mediated potassium efflux and hyperpolarization of the cell. K_ATP_ channels are inhibited by sulfonylureas, such as tolbutamide, preventing the protective effects of ketosis. Conversely, in the absence of ketosis, K_ATP_ channel openers, such as diazoxide, allow potassium efflux and recapitulate the protective effect of ketosis.

Our data also suggest that K_ATP_ channels are necessary for a ketogenic diet to provide analgesia. Adding tolbutamide could prevent the antinociceptive activity of a ketogenic diet in response to capsaicin injection. Our data agree with prior studies where treatment with tolbutamide is not pro-nociceptive and, by itself, does not cause allodynia or evoke nocifensive behaviors [45]. Tolbutamide is a specific inhibitor for K_ATP_ channels containing SUR1 subunits [3; 4], suggesting that SUR1-containing K_ATP_ channels are involved in ketogenic diet-mediated analgesia. Consistent with this view, activating K_ATP_ channels in chow-fed mice with diazoxide mimicked the effects of a ketogenic diet via reductions in pain behaviors and spinal expression of p-ERK. Diazoxide is also a selective agonist of SUR1-containing K_ATP_ channels, having no effect on SUR2A and minimally activating SUR2B-containing K_ATP_ channels [36; 47; 69]. Importantly, levcromakalim, a SUR2A and SUR2B-containing K_ATP_ channel agonist [36; 47; 59; 69], did not prevent capsaicin-evoked nociception in the mouse hind paw. We propose a ketogenic diet and ketone oxidation regulate SUR1-containing K_ATP_ channels that alter behavioral responses to a broad spectrum of noxious insults.

This mechanism may appear to be at odds with recent studies describing migraine induction following treatment with K_ATP_ channel activators [1; 13] and the ability of a ketogenic diet to reduce migraine [9; 19]. In these studies, migraines were induced in patients given subcutaneous levcromakalim, a specific agonist of SUR2A and SUR2B-containing K_ATP_ channels [36; 47; 59; 69]. This discrepancy may be related to the selective pharmacology between SUR1- and SUR2-isoform-containing K_ATP_ channel complexes. This view is supported by two pieces of data presented here: 1) tolbutamide, a specific antagonist of SUR1-containing K_ATP_ channels [3; 4], reduced the protective effect of a ketogenic diet against capsaicin, and 2) levcromakalim was unable to prevent capsaicin-evoked nociception or early activation in the spinal dorsal horn.

Additional supportive evidence comes from studies revealing interactions with K_ATP_ channel activity and opioid signaling [29; 41; 42; 53; 57]. SUR1 knockout or SUR1 inhibition reduces opioid-mediated analgesia [29; 53; 57], and diazoxide-mediated analgesia is partially reduced by knockdown or pharmacological inhibition of μ-, κ-, and δ-opioid receptors [41; 42]. Together with the data presented here, these findings suggest that a ketogenic diet may enhance opioid-mediated analgesia downstream of K_ATP_ channels. As intermittent fasting is associated with increased ketogenesis and ketone oxidation [10; 17; 51], this notion is consistent with reports that intermittent fasting enhances antinociception following morphine administration in mice [21]. Thus, it will be important to explore endogenous opioid production and opioid receptor signaling downstream of ketone oxidation and modulation of K_ATP_ channels in future studies.

Growing evidence suggests that ketogenic diets can play a role in preventing and reversing nociception in preclinical and clinical settings [9; 15; 19; 25; 26; 28; 55; 56; 73]. Unfortunately, these diets are restrictive to many, and adherence limits their use. Thus, pharmacologically mimicking the antinociception provided by a ketogenic diet is an attractive therapeutic strategy. Here, we demonstrate that a ketogenic diet provides antinociception to a range of noxious stimuli downstream of ketolysis in peripheral sensory neurons. Further, we demonstrate that a ketogenic diet requires K_ATP_ channel activity to mediate this protective effect and that activating K_ATP_ channels without ketosis is sufficient to mimic the antinociception provided by a ketogenic diet. This work suggests targeting SUR1-containing K_ATP_ channels could recapitulate the protective effects of a ketogenic diet without stringent dietary intervention.

## Acknowledgments

This work was supported by NIH grants R01 NS043314 (DEW), R01 AG069781 (PAC), R01 DK091538 (PAC), the Kansas Institutional Development Award (IDeA) P20 GM103418, Kansas University Training Program in Neurological and Rehabilitation Sciences NIH T32HD057850 (JE), Translating Obesity, Metabolic Dysfunction, and Comorbid Disease States NIH T32DK128770 (ST), the Kansas IDDRC P30HD00228, and University of Minnesota Institute for Diabetes, Obesity, and Metabolism.

## Conflict of interest

D.E.W. is conducting unrelated research under contract with Annexon Biosciences. P.A.C. has consulted for Pfizer, Inc., Abbott Laboratories, and Jansen Research & Development. The other authors declare no competing financial interests.

## Author contributions

JE and DEW designed the research study; JE, JMR, PL, JJ, and ST performed the experiments; JE and DEW analyzed the data; all authors contributed to the manuscript.

## Bibliography

[1] Al-Karagholi MA-M, Hansen JM, Guo S, Olesen J, Ashina M. Opening of ATP-sensitive potassium channels causes migraine attacks: a new target for the treatment of migraine. Brain 2019;142(9):2644–2654.

[2] Andersson DA, Gentry C, Light E, Vastani N, Vallortigara J, Bierhaus A, Fleming T, Bevan S. Methylglyoxal evokes pain by stimulating TRPA1. PloS one 2013;8(10):e77986.

[3] Ashfield R, Gribble FM, Ashcroft SJ, Ashcroft FM. Identification of the high-affinity tolbutamide site on the SUR1 subunit of the K(ATP) channel. Diabetes 1999;48(6):1341–1347.

[4] Babenko AP, Gonzalez G, Bryan J. The tolbutamide site of SUR1 and a mechanism for its functional coupling to KATP channel closure. FEBS Letters 1999;459(3):367–376.

[5] Barragán-Iglesias P, Kuhn J, Vidal-Cantú GC, Salinas-Abarca AB, Granados-Soto V, Dussor GO, Campbell ZT, Price TJ. Activation of the integrated stress response in nociceptors drives methylglyoxal-induced pain. Pain 2019;160(1):160.

[6] Bekircan-Kurt CE, Üçeyler N, Sommer C. Cutaneous activation of rage in nonsystemic vasculitic and diabetic neuropathy. Muscle & Nerve 2014;50(3):377–383.

[7] Bestall SM, Hulse RP, Blackley Z, Swift M, Ved N, Paton K, Beazley-Long N, Bates DO, Donaldson LF. Sensory neuronal sensitisation occurs through HMGB-1–RAGE and TRPV1 in high-glucose conditions. Journal of Cell Science 2018;131(14).

[8] Bierhaus A, Fleming T, Stoyanov S, Leffler A, Babes A, Neacsu C, Sauer SK, Eberhardt M, Schnölzer M, Lasitschka F. Methylglyoxal modification of Na v 1.8 facilitates nociceptive neuron firing and causes hyperalgesia in diabetic neuropathy. Nature medicine 2012;18(6):926–933.

[9] Bongiovanni D, Benedetto C, Corvisieri S, Del Favero C, Orlandi F, Allais G, Sinigaglia S, Fadda M. Effectiveness of ketogenic diet in treatment of patients with refractory chronic migraine. Neurological Sciences 2021;42(9):3865–3870.

[10] Cahill Jr GF. Fuel metabolism in starvation. Annu Rev Nutr 2006;26:1–22.

[11] Campanella ME, Chu H, Low PS. Assembly and regulation of a glycolytic enzyme complex on the human erythrocyte membrane. Proceedings of the National Academy of Sciences 2005;102(7):2402–2407.

[12] Chen Y, Ouyang X, Hoque R, Garcia-Martinez I, Yousaf MN, Tonack S, Offermanns S, Dubuquoy L, Louvet A, Mathurin P. β-Hydroxybutyrate protects from alcohol-induced liver injury via a Hcar2-cAMP dependent pathway. Journal of hepatology 2018;69(3):687–696.

[13] Christensen SL, Rasmussen RH, Ernstsen C, La Cour S, David A, Chaker J, Haanes KA, Christensen ST, Olesen J, Kristensen DM. CGRP-dependent signalling pathways involved in mouse models of GTN-cilostazol- and levcromakalim-induced migraine. Cephalalgia 2021;41(14):1413–1426.

[14] Cooper MA, McCoin C, Pei D, Thyfault JP, Koestler D, Wright DE. Reduced mitochondrial reactive oxygen species production in peripheral nerves of mice fed a ketogenic diet. Experimental physiology 2018;103(9):1206–1212.

[15] Cooper MA, Menta BW, Perez-Sanchez C, Jack MM, Khan ZW, Ryals JM, Winter M, Wright DE. A ketogenic diet reduces metabolic syndrome-induced allodynia and promotes peripheral nerve growth in mice. Experimental Neurology 2018;306:149–157.

[16] Coste B, Mathur J, Schmidt M, Earley TJ, Ranade S, Petrus MJ, Dubin AE, Patapoutian A. Piezo1 and Piezo2 are essential components of distinct mechanically activated cation channels. Science 2010;330(6000):55-60.

[17] Cotter DG, Schugar RC, Wentz AE, André d’Avignon D, Crawford PA. Successful adaptation to ketosis by mice with tissue-specific deficiency of ketone body oxidation. American Journal of Physiology-Endocrinology and Metabolism 2013;304(4):E363–E374.

[18] Dhar-Chowdhury P, Harrell MD, Han SY, Jankowska D, Parachuru L, Morrissey A, Srivastava S, Liu W, Malester B, Yoshida H, Coetzee WA. The glycolytic enzymes, glyceraldehyde-3-phosphate dehydrogenase, triose-phosphate isomerase, and pyruvate kinase are components of the K(ATP) channel macromolecular complex and regulate its function. J Biol Chem 2005;280(46):38464–38470.

[19] Di Lorenzo C, Coppola G, Bracaglia M, Di Lenola D, Sirianni G, Rossi P, Di Lorenzo G, Parisi V, Serrao M, Cervenka MC. A ketogenic diet normalizes interictal cortical but not subcortical responsivity in migraineurs. BMC neurology 2019;19(1):1–9.

[20] Düll MM, Riegel K, Tappenbeck J, Ries V, Strupf M, Fleming T, Sauer SK, Namer B. Methylglyoxal causes pain and hyperalgesia in human through C-fiber activation. Pain 2019;160(11):2497–2507.

[21] Duron DI, Hanak F, Streicher JM. Daily intermittent fasting in mice enhances morphine-induced antinociception while mitigating reward, tolerance, and constipation. PAIN 2020;161(10).

[22] Eberhardt MJ, Filipovic MR, Leffler A, de la Roche J, Kistner K, Fischer MJ, Fleming T, Zimmermann K, Ivanovic-Burmazovic I, Nawroth PP. Methylglyoxal activates nociceptors through transient receptor potential channel A1 (TRPA1): a possible mechanism of metabolic neuropathies. Journal of Biological Chemistry 2012;287(34):28291–28306.

[23] Edmond J, Robbins RA, Bergstrom JD, Cole RA, de Vellis J. Capacity for substrate utilization in oxidative metabolism by neurons, astrocytes, and oligodendrocytes from developing brain in primary culture. Journal of Neuroscience Research 1987;18(4):551–561.

[24] Enders J, Jack J, Thomas S, Lynch P, Lasnier S, Cao X, Swanson MT, Ryals JM, Thyfault JP, Puchalska P, Crawford PA, Wright DE. Ketolysis is required for the proper development and function of the somatosensory nervous system. Experimental Neurology 2023:114428.

[25] Enders J, Swanson MT, Ryals J, Wright DE. A ketogenic diet reduces mechanical allodynia and improves epidermal innervation in diabetic mice. Pain 2022;163(4):682–689.

[26] Enders J, Thomas S, Swanson MT, Ryals JM, Wright DE. A ketogenic diet prevents methylglyoxal-evoked nociception by scavenging methylglyoxal. J Pain 2022;23(5):24.

[27] Epstein T, Xu L, Gillies RJ, Gatenby RA. Separation of metabolic supply and demand: aerobic glycolysis as a normal physiological response to fluctuating energetic demands in the membrane. Cancer & Metabolism 2014;2(1):7.

[28] Field R, Pourkazemi F, Rooney K. Effects of a Low-Carbohydrate Ketogenic Diet on Reported Pain, Blood Biomarkers and Quality of Life in Patients with Chronic Pain: A Pilot Randomized Clinical Trial. Pain Medicine 2022;23(2):326–338.

[29] Fisher C, Johnson K, Okerman T, Jurgenson T, Nickell A, Salo E, Moore M, Doucette A, Bjork J, Klein AH. Morphine Efficacy, Tolerance, and Hypersensitivity Are Altered After Modulation of SUR1 Subtype KATP Channel Activity in Mice. Frontiers in Neuroscience 2019;13.

[30] Gribble FM, Tucker SJ, Ashcroft FM. The essential role of the Walker A motifs of SUR1 in K-ATP channel activation by Mg-ADP and diazoxide. The EMBO Journal 1997;16(6):1145–1152.

[31] Griggs RB, Laird DE, Donahue RR, Fu W, Taylor BK. Methylglyoxal requires AC1 and TRPA1 to produce pain and spinal neuron activation. Frontiers in neuroscience 2017;11:679.

[32] Griggs RB, Santos DF, Laird DE, Doolen S, Donahue RR, Wessel CR, Fu W, Sinha GP, Wang P, Zhou J. Methylglyoxal and a spinal TRPA1-AC1-Epac cascade facilitate pain in the db/db mouse model of type 2 diabetes. Neurobiology of disease 2019;127:76–86.

[33] Hiyama H, Yano Y, So K, Imai S, Nagayasu K, Shirakawa H, Nakagawa T, Kaneko S. TRPA1 sensitization during diabetic vascular impairment contributes to cold hypersensitivity in a mouse model of painful diabetic peripheral neuropathy. Molecular pain 2018;14:1744806918789812–1744806918789812.

[34] Ho T, Potapenko E, Davis DB, Merrins MJ. A plasma membrane-associated glycolytic metabolon is functionally coupled to K_ATP_ channels in pancreatic α and β cells from humans and mice. Cell Reports 2023;42(4).

[35] Huang Q, Chen Y, Gong N, Wang Y-X. Methylglyoxal mediates streptozotocin-induced diabetic neuropathic pain via activation of the peripheral TRPA1 and Nav1. 8 channels. Metabolism 2016;65(4):463-474.

[36] Inagaki N, Gonoi T, Iv JPC, Wang C-Z, Aguilar-Bryan L, Bryan J, Seino S. A Family of Sulfonylurea Receptors Determines the Pharmacological Properties of ATP-Sensitive K+ Channels. Neuron 1996;16(5):1011–1017.

[37] Kawamura M, Ruskin DN, Masino SA. Metabolic autocrine regulation of neurons involves cooperation among pannexin hemichannels, adenosine receptors, and KATP channels. Journal of Neuroscience 2010;30(11):3886–3895.

[38] Kawano T, Zoga V, McCallum JB, Wu HE, Gemes G, Liang MY, Abram S, Kwok WM, Hogan QH, Sarantopoulos CD. ATP-sensitive potassium currents in rat primary afferent neurons: biophysical, pharmacological properties, and alterations by painful nerve injury. Neuroscience 2009;162(2):431–443.

[39] Lam D, Momeni Z, Theaker M, Jagadeeshan S, Yamamoto Y, Ianowski JP, Campanucci VA. RAGE-dependent potentiation of TRPV1 currents in sensory neurons exposed to high glucose. PLOS ONE 2018;13(2):e0193312.

[40] Lee W, Leddy HA, Chen Y, Lee SH, Zelenski NA, McNulty AL, Wu J, Beicker KN, Coles J, Zauscher S, Grandl J, Sachs F, Guilak F, Liedtke WB. Synergy between Piezo1 and Piezo2 channels confers high-strain mechanosensitivity to articular cartilage. Proceedings of the National Academy of Sciences 2014;111(47):E5114–E5122.

[41] Lohmann AB, Welch SP. Antisenses to opioid receptors attenuate ATP-gated K+ channel opener-induced antinociception. European Journal of Pharmacology 1999;384(2):147–152.

[42] Lohmann AB, Welch SP. ATP-gated K+ channel openers enhance opioid antinociception: indirect evidence for the release of endogenous opioid peptides. European Journal of Pharmacology 1999;385(2):119–127.

[43] Lu Y, Yang Y-Y, Zhou M-W, Liu N, Xing H-Y, Liu X-X, Li F. Ketogenic diet attenuates oxidative stress and inflammation after spinal cord injury by activating Nrf2 and suppressing the NF-κB signaling pathways. Neuroscience Letters 2018;683:13–18.

[44] Lund TM, Ploug KB, Iversen A, Jensen AA, Jansen-Olesen I. The metabolic impact of β-hydroxybutyrate on neurotransmission: Reduced glycolysis mediates changes in calcium responses and KATP channel receptor sensitivity. J Neurochem 2015;132(5):520–531.

[45] Luu W, Bjork J, Salo E, Entenmann N, Jurgenson T, Fisher C, Klein AH. Modulation of SUR1 K(ATP) Channel Subunit Activity in the Peripheral Nervous System Reduces Mechanical Hyperalgesia after Nerve Injury in Mice. Int J Mol Sci 2019;20(9).

[46] Ma W, Berg J, Yellen G. Ketogenic diet metabolites reduce firing in central neurons by opening KATP channels. Journal of Neuroscience 2007;27(14):3618–3625.

[47] Matsuoka T, Matsushita K, Katayama Y, Fujita A, Inageda K, Tanemoto M, Inanobe A, Yamashita S, Matsuzawa Y, Kurachi Y. C-Terminal Tails of Sulfonylurea Receptors Control ADP-Induced Activation and Diazoxide Modulation of ATP-Sensitive K^+^ Channels. Circulation Research 2000;87(10):873–880.

[48] Mikesell AR, Isaeva O, Moehring F, Sadler KE, Menzel AD, Stucky CL. Keratinocyte PIEZO1 modulates cutaneous mechanosensation. Elife 2022;11:e65987.

[49] Nakai-Shimoda H, Himeno T, Okawa T, Miura-Yura E, Sasajima S, Kato M, Yamada Y, Morishita Y, Tsunekawa S, Kato Y. Kir6. 2-deficient mice develop somatosensory dysfunction and axonal loss in the peripheral nerves. Iscience 2022;25(1):103609.

[50] Pitcher MH, Von Korff M, Bushnell MC, Porter L. Prevalence and Profile of High-Impact Chronic Pain in the United States. J Pain 2019;20(2):146–160.

[51] Puchalska P, Nelson AB, Stagg DB, Crawford PA. Determination of ketone bodies in biological samples via rapid UPLC-MS/MS. Talanta 2021;225:122048.

[52] Puljung M, Vedovato N, Usher S, Ashcroft F. Activation mechanism of ATP-sensitive K+ channels explored with real-time nucleotide binding. eLife 2019;8:e41103.

[53] Rodrigues AR, Duarte ID. The peripheral antinociceptive effect induced by morphine is associated with ATP-sensitive K(+) channels. Br J Pharmacol 2000;129(1):110–114.

[54] Rojas DR, Tegeder I, Kuner R, Agarwal N. Hypoxia-inducible factor 1α protects peripheral sensory neurons from diabetic peripheral neuropathy by suppressing accumulation of reactive oxygen species. Journal of Molecular Medicine 2018;96(12):1395–1405.

[55] Ruskin DN, Kawamura Jr M, Masino SA. Reduced pain and inflammation in juvenile and adult rats fed a ketogenic diet. PloS one 2009;4(12):e8349.

[56] Ruskin DN, Sturdevant IC, Wyss LS, Masino SA. Ketogenic diet effects on inflammatory allodynia and ongoing pain in rodents. Scientific Reports 2021;11(1):1–8.

[57] Sakamaki G, Johnson K, Mensinger M, Hmu E, Klein AH. Loss of SUR1 subtype KATP channels alters antinociception and locomotor activity after opioid administration. Behavioural Brain Research 2021;414:113467.

[58] Salomón T, Sibbersen C, Hansen J, Britz D, Svart MV, Voss TS, Møller N, Gregersen N, Jørgensen KA, Palmfeldt J, Poulsen TB, Johannsen M. Ketone Body Acetoacetate Buffers Methylglyoxal via a Non-enzymatic Conversion during Diabetic and Dietary Ketosis. Cell Chemical Biology 2017;24(8):935–943.e937.

[59] Schwanstecher M, Sieverding C, Dörschner H, Gross I, Aguilar-Bryan L, Schwanstecher C, Bryan J. Potassium channel openers require ATP to bind to and act through sulfonylurea receptors. The EMBO Journal 1998;17(19):5529–5535.

[60] Shang S, Wang L, Zhang Y, Lu H, Lu X. The beta-hydroxybutyrate suppresses the migration of glioma cells by inhibition of NLRP3 inflammasome. Cellular and molecular neurobiology 2018;38(8):1479–1489.

[61] Staruschenko A, Jeske NA, Akopian AN. Contribution of TRPV1-TRPA1 interaction to the single channel properties of the TRPA1 channel. Journal of biological chemistry 2010;285(20):15167–15177.

[62] Tucker SJ, Gribble FM, Proks P, Trapp S, Ryder TJ, Haug T, Reimann F, Ashcroft FM. Molecular determinants of KATP channel inhibition by ATP. The EMBO Journal 1998;17(12):3290–3296.

[63] Valdebenito R, Ruminot I, Garrido-Gerter P, Fernández-Moncada I, Forero-Quintero L, Alegría K, Becker HM, Deitmer JW, Barros LF. Targeting of astrocytic glucose metabolism by beta-hydroxybutyrate. Journal of Cerebral Blood Flow & Metabolism 2015;36(10):1813–1822.

[64] Wang J, La J-H, Hamill OP. PIEZO1 is selectively expressed in small diameter mouse DRG neurons distinct from neurons strongly expressing TRPV1. Frontiers in Molecular Neuroscience 2019;12:178.

[65] Wang M, Wu J-X, Ding D, Chen L. Structural insights into the mechanism of pancreatic KATP channel regulation by nucleotides. Nature Communications 2022;13(1):2770.

[66] Wang X, Wu X, Liu Q, Kong G, Zhou J, Jiang J, Wu X, Huang Z, Su W, Zhu Q. Ketogenic Metabolism Inhibits Histone Deacetylase (HDAC) and Reduces Oxidative Stress After Spinal Cord Injury in Rats. Neuroscience 2017;366:36–43.

[67] Wei J-Y, Liu C-C, Ouyang H-D, Ma C, Xie M-X, Liu M, Lei W-L, Ding H-H, Wu S-L, Xin W-J. Activation of RAGE/STAT3 pathway by methylglyoxal contributes to spinal central sensitization and persistent pain induced by bortezomib. Experimental neurology 2017;296:74–82.

[68] Weng H-J, Patel KN, Jeske NA, Bierbower SM, Zou W, Tiwari V, Zheng Q, Tang Z, Mo GC, Wang Y. Tmem100 is a regulator of TRPA1-TRPV1 complex and contributes to persistent pain. Neuron 2015;85(4):833–846.

[69] Wheeler A, Wang C, Yang K, Fang K, Davis K, Styer AM, Mirshahi U, Moreau C, Revilloud J, Vivaudou M, Liu S, Mirshahi T, Chan KW. Coassembly of different sulfonylurea receptor subtypes extends the phenotypic diversity of ATP-sensitive potassium (KATP) channels. Mol Pharmacol 2008;74(5):1333–1344.

[70] Yong RJ, Mullins PM, Bhattacharyya N. Prevalence of chronic pain among adults in the United States. PAIN 2022;163(2).

[71] Youm Y-H, Nguyen KY, Grant RW, Goldberg EL, Bodogai M, Kim D, D’agostino D, Planavsky N, Lupfer C, Kanneganti TD. The ketone metabolite β-hydroxybutyrate blocks NLRP3 inflammasome– mediated inflammatory disease. Nature medicine 2015;21(3):263–269.

[72] Zherebitskaya E, Akude E, Smith DR, Fernyhough P. Development of Selective Axonopathy in Adult Sensory Neurons Isolated From Diabetic Rats: Role of Glucose-Induced Oxidative Stress. Diabetes 2009;58(6):1356–1364.

[73] Zhong S, Zhou Z, Lin X, Liu F, Liu C, Liu Z, Deng W, Zhang X, Chang H, Zhao C. Ketogenic diet prevents paclitaxel-induced neuropathic nociception through activation of PPARγ signalling pathway and inhibition of neuroinflammation in rat dorsal root ganglion. European Journal of Neuroscience 2021;54(4):5341–5356.

